# Aimed Limb Movements in a Hemimetabolous Insect are Intrinsically Compensated for Allometric Wing Growth by Developmental Mechanisms

**DOI:** 10.1101/662759

**Authors:** Alexandra J. Patel, Thomas Matheson

## Abstract

For aimed limb movements to remain functional they must be adapted to developmental changes in body morphology and sensory-motor systems. Insects use their limbs to groom the body surface or to dislodge external stimuli, but they face the particular problem of adapting these movements to step-like changes in body morphology during metamorphosis or moulting. Locusts are hemimetabolous insects in which the imaginal moult to adulthood results in a sudden and dramatic allometric growth of the wings relative to the body and the legs. We show that, despite this, hind limb scratches aimed at mechanosensory stimuli on the wings remain targeted to appropriate locations after moulting. In juveniles, the tips of the wings extend less than half way along the abdomen, but in adults they extend well beyond the posterior end. Kinematic analyses were used to examine the scratching responses of juveniles (5^th^ instars) and adults to touch of anterior (wing base) and posterior (distal abdomen) targets that develop isometrically, and to wing tip targets that are anterior in juveniles but posterior in adults. Juveniles reach the (anterior) wing tip with the distal tibia of the hind leg using anterior rotation of the thoraco-coxal and coxo-trochanteral (‘hip’) joints and flexion of the femoro-tibial (‘knee’) joint. Adults, however, reach the corresponding (but now posterior) wing tip using posterior rotation of the hip and extension of the knee, reflecting a different underlying motor pattern. This change in kinematics occurs immediately after the adult moult without learning, indicating that the switch is developmentally programmed.

**SUMMARY STATEMENT:** A developmentally programmed change in the scratching movements of locusts permits adult animals to aim their movements at new wing tip targets, without learning.

## INTRODUCTION

Fundamentally important behaviours such as reaching, stepping, grooming and prey capture all require accurately aimed limb movements. The neuronal networks underlying these movements must be adapted to changes in body morphology during development so that they remain functional throughout life. Sensory-motor plasticity is thus critical for tuning and maintaining behaviours during development (Easter, 1983). For example, human infants begin reaching towards targets at age 4-5 months, but the kinematics of their movements change over the first three years of life before assuming their adult-like patterns (Konczak and Dichgans, 1997). This reflects continual recalibration of the sensory-motor system as both the peripheral mechanics and the central nervous system develop (Konczak *et al.* 1997). Adult locusts spontaneously walk across larger gaps than juveniles, apparently judging the distance using visual cues. The change in maximum gap width crossed is reportedly not expressed immediately after the moult, but becomes apparent with some delay; and appears to be facilitated by a period of interaction with a complex external environment (Ben-Nun *et al.* 2013) – again suggesting age-dependent recalibration. Praying mantises that capture prey using visually triggered strikes of their raptorial front legs have longer strike distances as adults than as juveniles. Striking is triggered at age- and body-size-appropriate distances largely as a result of concomitant changes in the size and shape of the eyes and the size of the body and legs at each moult (Balderrama and Maldonado, 1973; Maldonado *et al.* 1974; Kral and Poteser, 2009) – so here it seems that experience-dependent ‘recalibration’ is not required.

In the case of grooming movements aimed at a target elsewhere on the body, differential growth of body and limbs can cause a change in the spatial relationship between the target and the grooming limb. Changes in the sensory system, such as addition or loss of exteroceptors or changes in their properties, may lead to changes in the representation of both the body surface and features of the external environment (Murphey *et al.* 1980; Chiba *et al.* 1988). Growth leads to changes in limb lengths and musculoskeletal properties which, given an unchanged motor pattern, would result in different and inappropriate movement trajectories. If behaviours are to be conserved during growth and development, then plasticity of the underlying neuronal pathways needs to match the biomechanical changes.

In locust scratching, deflection of tactile hairs on a wing elicits movement of the ipsilateral hind leg, with the distal end of the tibia and tarsus aimed toward the site of stimulation (Berkowitz and Laurent, 1996a; Matheson, 1997, 1998; Dürr and Matheson, 2003).

Furthermore, stimulation of different locations along the wing surface elicits tarsal trajectories that are target-specific from onset and accurately aimed to each stimulus site (Berkowitz and Laurent, 1996a; Matheson, 1997, surface of the hind wing that is outermost1998; Dürr and Matheson, 2003). To reach progressively more proximal targets requires increases in the elevation of the femur and increased flexion of the femoro-tibial joint (Fig. 1). These continuously graded changes in hind leg joint angles suggest that adult locusts use a single, systematically varied, movement pattern for scratching, rather than using distinct motor strategies specific to each location within the wing receptive field (Dürr and Matheson, 2003). In addition, locusts compensate their scratching movements against perturbations caused by loading the hind leg (Matheson and Dürr, 2003). This plastic behaviour provides an opportunity to investigate how aimed movements adapt to morphological changes that arise during development.

**Figure 1.**
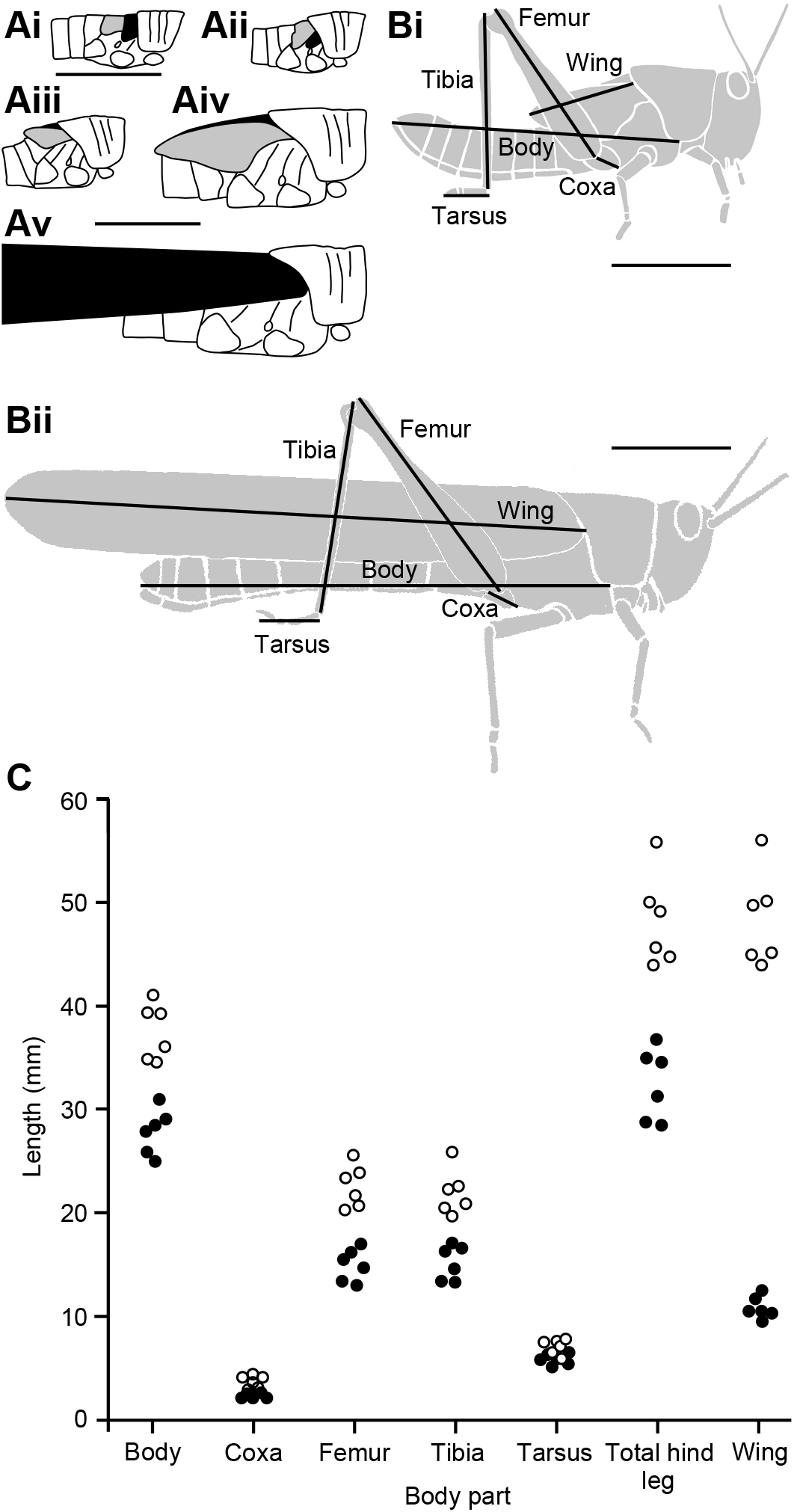
Comparison of body and limb dimensions for the 5^th^ instar and adult *Schistocerca gregaria* used in subsequent experiments. A. Schematic representation of wing bud rotation during development. Each diagram illustrates the pronotum, thorax and first three abdominal segments, with the fore wing shaded black and the hind wing shaded grey. Ai is reproduced at 2× scale compared to Aii-v; scale bar 5 mm for Ai, 10 mm for Aii-Av). Ai. 2^nd^ instar: the wing buds are oriented with future leading edges anterio-ventral, dorsal surfaces facing outwards. Aii. 3^rd^ instar: The hind wing bud partially covers the fore wing bud, orientations unchanged. Aiii. 4^th^ instar: both wing buds have rotated upwards so that their future leading edges are oriented dorsally, ventral surfaces of the wings facing outwards. Aiv. 5^th^ instar: orientations unchanged – the hind wing bud covers the fore wing bud, with its ventral surface facing outwards. Av. Adult: the much enlarged adult wings have rotated back to their original positions, with the leading edges ventral. The fore wing (black) covers the folded hind wing (not visible), with its dorsal surface facing outwards. B. Dimensions measured for 5^th^ instars (Bi) and adults (Bii). Both scale bars 10 mm. C. Measured values of each dimension and summed total hind leg length (coxa + femur + tibia), for six 5^th^ instars and six different adults. Fore wing length increased disproportionately more than total hind leg length. See Table 1 for a summary.

Hemimetabolous insects like locusts maintain essentially the same body structure throughout life, enlarging abruptly – but relatively proportionately – at each successive moult (Uvarov, 1966). At the adult (imaginal) moult, however, wing length increases disproportionately relative to body and leg length. This allometric growth means that the wing tips lie relatively much further posterior in adults than in the last juvenile stage (5^th^ instar). To remain functional, scratching movements elicited by tactile stimulation of the wings must therefore adapt to the relative changes in limb and wing dimensions. To complicate matters further, there are also two morphological rotations of the wings during development (Roonwal, 1940, 1946; Ivanova, 1947; Burnett, 1951; Uvarov, 1966; Dirsh, 1967; Khattar, 1972). In *Schistocerca gregaria*, wing pads develop in 1^st^ instars as ventrally directed extensions of the dorsal cuticular plates. The fore wing pad lies anterior to the hind wing pad, and both have their dorsal surfaces outermost. The wing pads remain in the same position in 2^nd^ and 3^rd^ instars (Fig. 1Ai,ii), but during the moult to 4^th^ instar both wing pads rotate: the ventral surfaces turn outermost, and the hind wing pad now largely covers the fore wing pad (Fig. 1Aiii; Roonwal, 1940). In 5^th^ instars the wing pads remain in the same position (Fig. 1Aiv), but on moulting to adulthood, they rotate again so that the dorsal surfaces of both wings are outermost, and the fore wing now covers the hind wing at rest (Fig. 1Av; Roonwal, 1952). In adults the dorsal surface of the fore wing is outermost, whereas in 5^th^ instars this surface is innermost and apposed to the abdomen. In 5^th^ instars it is the ventral surface of the hind wing that is outermost (Fig. 1Av *vs* 1Aiv).

Tactile stimuli that elicit aimed scratching movements are detected by wing mechanosensory hairs (Page and Matheson, 2004). Developmental rotation of the wings at the imaginal moult means that those hairs which elicit scratching in resting 5^th^ instars and adults are necessarily located on different wing surfaces. In 5^th^ instars (where it is the ventral hind wing that is exposed) the sensory pathway originates from metathoracic wing hairs and passes to the metathoracic ganglion where the interneurons and motor neurons driving leg movement are located (Matheson, 1997). In adults, the sensory pathway originates from mesothoracic wing hairs on the exposed dorsal fore wing surface (Page and Matheson, 2004) and passes via the mesothoracic ganglion (Campbell, 1961) and descending connectives to eventually drive the same metathoracic motor networks (Matheson, 2002). At the imaginal moult locusts must therefore adapt their scratching movements to not only account for allometric elongation of the wing sensory surface, but also for a change from receptors on one surface to those on another.

Here we describe the scratching movements of 5^th^ instar locusts and show that for similar targets they have similar kinematics to those of adults. Both 5^th^ instar and adult locusts accurately aim their hind limb movements at distal wing tip targets despite the profound developmental changes that occur at the imaginal moult. The compensation for wing allometry and developmental rotation is apparent immediately after the moult, indicating that the animal’s internal body image is adjusted by developmental rather than learned mechanisms.

## RESULTS

### Wings grow allometrically at the imaginal moult

Body, leg and wing lengths were measured in 6 animals prior to and 6 different animals following the imaginal moult (Fig. 1Bi,ii, 1C). Adult bodies were 135% longer than those of 5^th^ instars, while hind leg segments were 122-161% longer (Table 1). In contrast, adult wings were 447% longer than the wing buds of 5^th^ instars (Fig. 1C, Table 1). Relative to the increase in body length, all hind limb segments thus increased isometrically by a factor of 0.9-1.2, whereas wing length increased allometrically by a factor of 3.3 (Table 1). The tips of the 5^th^ instar wing pads lay approximately 15 mm anterior to the tip of the abdomen, whereas the tips of the adult wings lay approximately 12 mm *posterior* to the tip of the abdomen (Fig. 1Bi,ii).

**Table 1.**
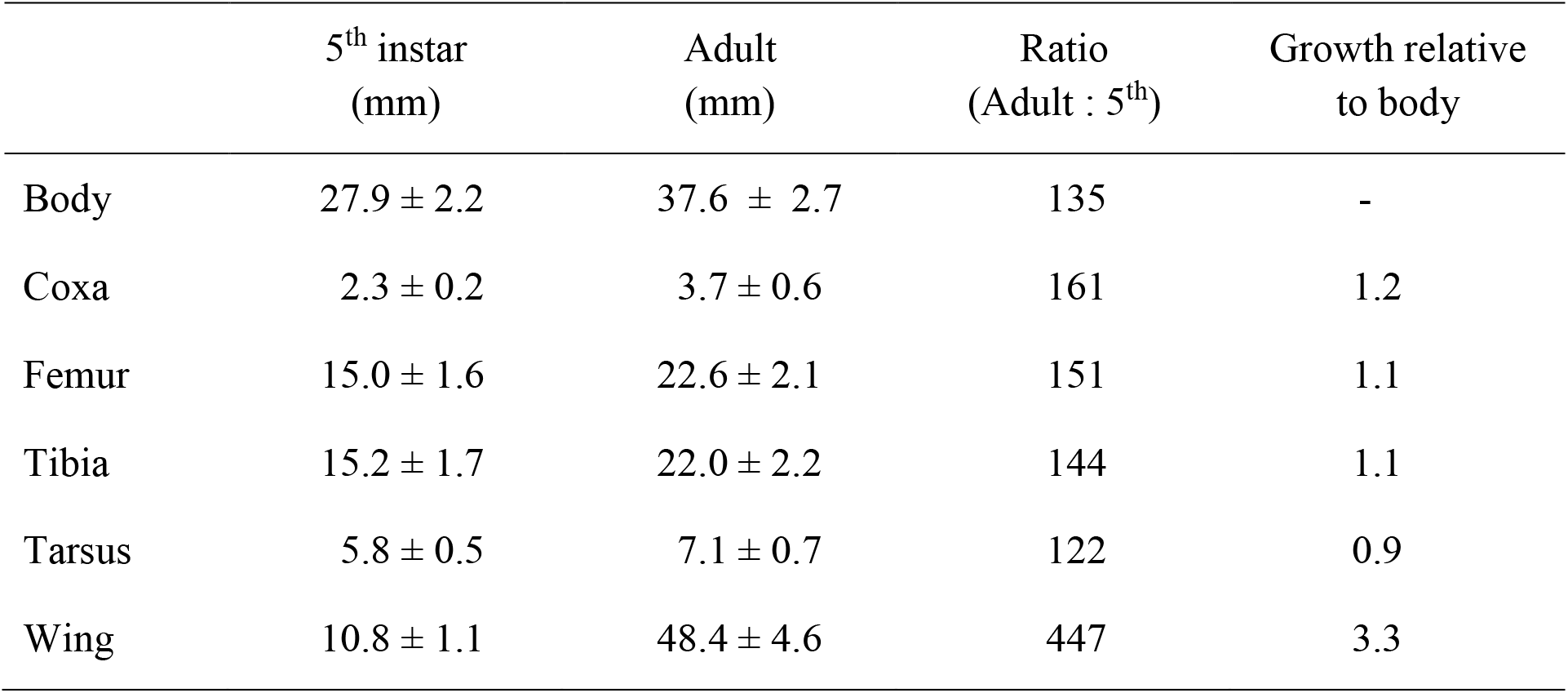
Comparison of body and limb dimensions for the 5^th^ instar and adult *Schistocerca gregaria* used in subsequent experiments. Values are means ± SD for six 5^th^ instars and six different adults. See Fig. 1 for the data and illustration of measurements.

|  | 5<sup>th</sup> instar<br>(mm) | Adult<br>(mm) | Ratio<br>(Adult : 5<sup>th</sup>) | Growth relative<br>to body |
| --- | --- | --- | --- | --- |
| Body | 27.9 ± 2.2 | 37.6 ± 2.7 | 135 | - |
| Coxa | 2.3 ± 0.2 | 3.7 ± 0.6 | 161 | 1.2 |
| Femur | 15.0 ± 1.6 | 22.6 ± 2.1 | 151 | 1.1 |
| Tibia | 15.2 ± 1.7 | 22.0 ± 2.2 | 144 | 1.1 |
| Tarsus | 5.8 ± 0.5 | 7.1 ± 0.7 | 122 | 0.9 |
| Wing | 10.8 ± 1.1 | 48.4 ± 4.6 | 447 | 3.3 |

### Scratching movement trajectories of 5^th^ instar and adult locusts

In resting adult locusts the dorsal fore wing surface is outermost and thus exposed. As a consequence of wing bud rotation during development, however, it is the ventral hind wing surface that is exposed in resting 5^th^ instars. Both 5^th^ instar and adult locusts performed scratching movements in response to touches on the exposed wing surfaces or abdomen (Fig. 2). Following stimulation of an anterior target at the base of the wing (yellow squares in Fig. 2Ai, Bi), 5^th^ instars, like adults, moved the distal end of the ipsilateral hind leg towards the target in an anteriorly-directed arc, and performed cyclical movements near to the target (black trajectories in Fig. 2Ai, Bi). Following stimulation of a posterior target on the tip of the abdomen (black squares in Fig. 2Aii, Bii) 5^th^ instars, again like adults, moved the distal end of the leg towards the target in posteriorly-directed movements and made cyclical movements near the target (black trajectories in Fig. 2Aii, Bii).

**Figure 2.**
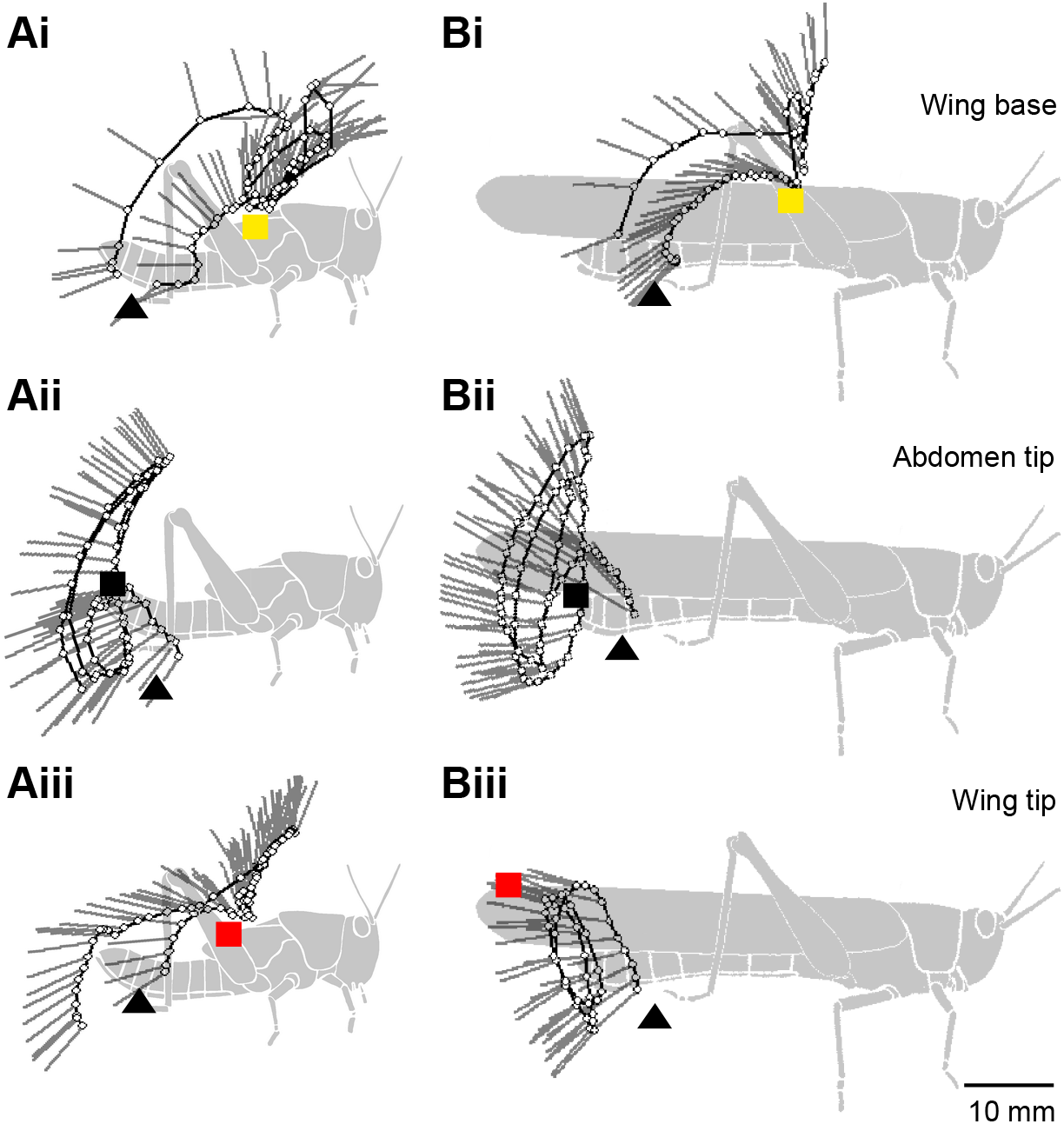
Both 5^th^ instar and adult locusts aim their scratching movements at targets on the wings and abdomen. Panels show trajectories of the distal tibia toward three different targets in 5^th^ instar (Ai-iii) and adult (Bi-iii) locusts. Stimulus locations on the wing base (Ai, Bi) are represented by yellow squares, those on the abdomen tip (Aii, Bii) by black squares, and those on the wing tip (Aiii, Biii) by red squares. Start positions are represented by black triangles. The positions of the tibio-tarsal joint (small open circles joined by black lines) and the tarsus (grey lines) are shown at intervals of 20 ms for each movement.

The hind wing tip in 5^th^ instars is an anterior target and its stimulation elicited anteriorly aimed movements (Fig. 2Aiii) which were similar in form to those aimed toward the hind wing base (Fig. 2Ai). In adults, however, the fore wing tip is a posterior target, and its stimulation elicited movements that were directed posteriorly (Fig. 2Biii). They were similar in form to those aimed towards nearby abdomen tip targets (Fig. 2Bii).

In 5^th^ instars, tactile stimulation of the normally hidden dorsal fore wing tip (see Methods) elicited anteriorly directed scratches. Of 110 trials in 11 locusts in which the fore wing tip was stimulated, 33 responses were aimed anterior to the midpoint of the abdomen, 19 were aimed anterior to the tip of the abdomen, and none were aimed posterior to the tip of the abdomen. The remaining 58 stimuli did not elicit scratching responses. Touching the dorsal fore wing tip therefore elicits anteriorly directed scratches in 5^th^ instars but posteriorly directed scratches in adults.

### Rapid change in behaviour at the imaginal moult

The switch from making anteriorly directed to posteriorly directed scratches in response to touch of the wing tip occurs instantaneously following the imaginal moult, and does not require a period of calibration or learning. In 9 of 9 animals observed continuously during moulting, the first touch of the newly expanded adult fore wing tip led immediately to posterior scratches, prior to any other significant hind limb movements, including walking or scratching. During the early part of moulting, animals make small wriggling movements while hanging head downwards to pull themselves out of their old exoskeleton, assisted by gravity. As soon as the adult emerges from the exuvium it rotates to a head up posture and remains essentially motionless while haemolymph is pumped through the wings to expand them, again assisted by gravity. These newly moulted animals responded reliably to tactile stimulation. To confirm that these posterior movements were retained beyond the moulting period, the hind legs of a further 13 animals – also observed continuously during moulting – were restrained immediately following the moult to prevent tibial extension that is required to reach the adult wing tip. On removal of the restraint 20-24h after the moult, all 13 of these animals responded immediately to touch of the adult fore wing tip with posterior scratches. In no case did the initial touch of the adult fore wing tip lead to an anterior scratch.

### Both 5^th^ instars and adults aim the distal tibia at wing and distal abdomen targets

For movements toward anterior wing base targets in both 5^th^ instars and adults, two different regions of the hind leg approach the target closely, resulting in a “w” shaped distribution of points of closest approach (Fig. 3Ai, Bi). Unit 4 at the centre of the femur (see Methods for a description of the division of the leg into analysis ‘units’) was closest for both 5^th^ instars and adults (double obelisk [‡] in Fig. 3Ai, Bi). A second minimum was centred on unit 18 of the distal tibia (single obelisk [†] in Fig. 3Ai, Bi). The physical constraints of hind leg movement mean that for the distal tibia or tarsus to reach the anterior targets (wing base and wing tip in 5^th^ instars; wing base in adults), the femur must rotate far anteriorly, itself crossing the target site (see Fig. 7Ci). The femur typically crossed the target once, and was then held anteriorly for the rest of the movement. In contrast, the distal tibia approached the target several times during the cyclic component of a scratch (Fig. 2). The local minimum at unit 18 thus reflected the aimed effector (distal tibia) whereas that at unit 4 reflected a mechanical consequence of that aimed movement. In movements toward (posterior) abdomen tip targets in both 5^th^ instars and adults, units 18 or 19 of the distal tibia most closely approached the target (single obelisks in Fig. 3Aii, Bii). The wing tip is an anterior target in 5^th^ instars, but a posterior target in adults, and this difference in target position is reflected in the parts of the hind leg that approach the target. In 5^th^ instars both unit 5 of the femur, and unit 19 of the distal tibia approached the target as described above for wing base targets (double obelisk and single obelisk, respectively in Fig. 3Aiii). In adults, only unit 23 of the tarsus reached the very posterior position of the wing tip target (single obelisk in Fig. 3Biii). The distal tibia and tarsus are thus used as effectors during scratching movements in both 5^th^ instars and adults, for both anterior and posterior targets.

**Figure 3.**
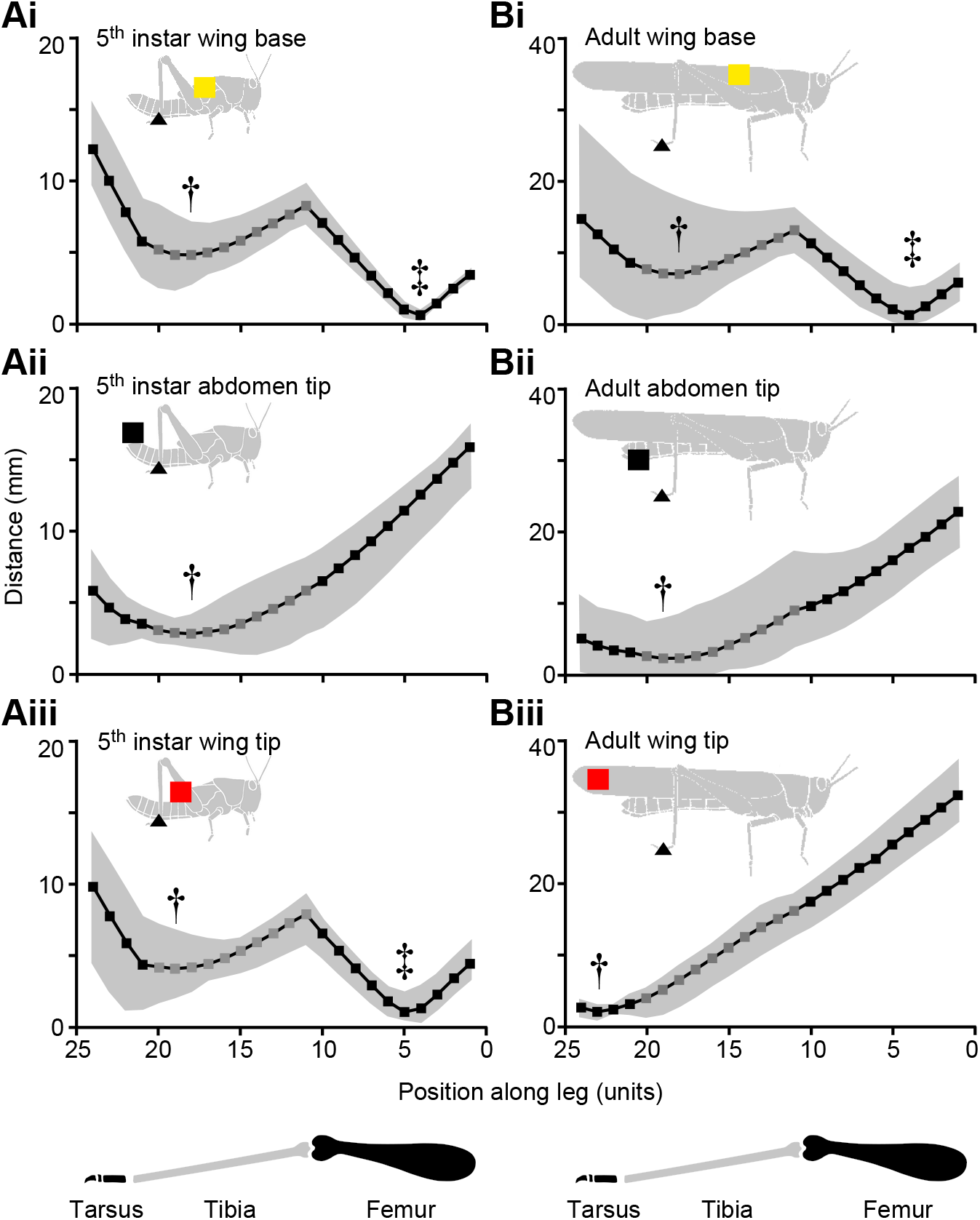
Both 5^th^ instar and adult locusts aim the distal end of the tibia at targets on the wings and abdomen. Lines and symbols show the mean closest approach of each leg ‘unit’ to the target during movements toward wing base (Ai, Bi, yellow squares), abdomen tip (Aii, Bii, black squares), and wing tip (Aiii, Biii, red squares) targets, for 5^th^ instar (A) and adult (B) locusts. Grey shading indicates the range. Dark grey symbols correspond to units of the tibia, black symbols correspond to units of the femur or tarsus (see inset diagrams at the bottom, and Methods). The region of the leg that approached the target most closely is labelled with a single obelisk (†) for the meaningful distal minima and a double obelisk (‡) for the femoral minima (see text). Note that the ordinate scales for 5^th^ instars and adults differ by a factor of two.

### Targeting accuracy is similar for all sites

The minimal distance from the distal effector point to the target during each trajectory provided a measure of accuracy. For 5^th^ instars, the mean minimal distance was similar for movements toward the wing base, abdomen tip and wing tip (repeated measures ANOVA, Greenhouse-Geisser correction, *F*_1.084,5.421_ = 0.448, *P* = 0.548). The mean minimal distances were 5.1 ± 0.8 mm, 5.1 ± 1.4 mm, and 4.3 ± 0.6 mm (mean ± SEM) for wing base, abdomen tip and wing tip targets respectively.

For adults, there was weak evidence that mean minimal distance may be different for different targets (repeated measures ANOVA, *F*_2, 10_ = 4.167, *P* = 0.048). There was no evidence for a difference in mean minimal distance for adult wing base and wing tip targets (posthoc two tailed *t*-test *P* = 0.537), and only weak evidence for differences between the abdomen tip target and either the wing base target (*P* = 0.028) or the wing tip target (*P* = 0.036). The mean minimal distances were 7.4 ± 1.7 mm, 3.5 ± 0.7 mm, and 6.2 ± 0.7 mm for wing base, abdomen tip and wing tip targets respectively.

There was no evidence for a difference between 5^th^ instars and adults in mean minimal distance for movements towards wing base targets (one way ANOVA, *F*_1, 10_ = 1.498, *P* = 0.249), abdomen tip targets (*F*_1, 10_ = 1.242, *P* = 0.291), or wing tip targets (*F*_1, 10_ = 3.815, *P* = 0.079). Overall, there is therefore little evidence for any difference in accuracy between sites or between ages.

### Initial direction of movement and speed is influenced by target site

The first 200 ms of each scratch was used to calculate a movement vector that was used to determine whether movements were target specific from their onset and thus predetermined. For 5^th^ instars, the initial directions of movement towards all three targets were directed dorsally and anteriorly (yellow, red and black vectors in Fig. 4A). There was no detectable effect of target position on the initial direction of movement, despite the fact that the abdomen tip target was posterior to the initial tarsus position while the two wing targets were anterior (non-parametric repeated measures ANOVA (Friedman Test): *χ*^2^_2,6_ = 0.333, *P* = 0.846). Nevertheless, the movement vectors were all correctly biased towards the appropriate targets (corresponding yellow, red and black squares in Fig. 4A inset).

**Figure 4.**
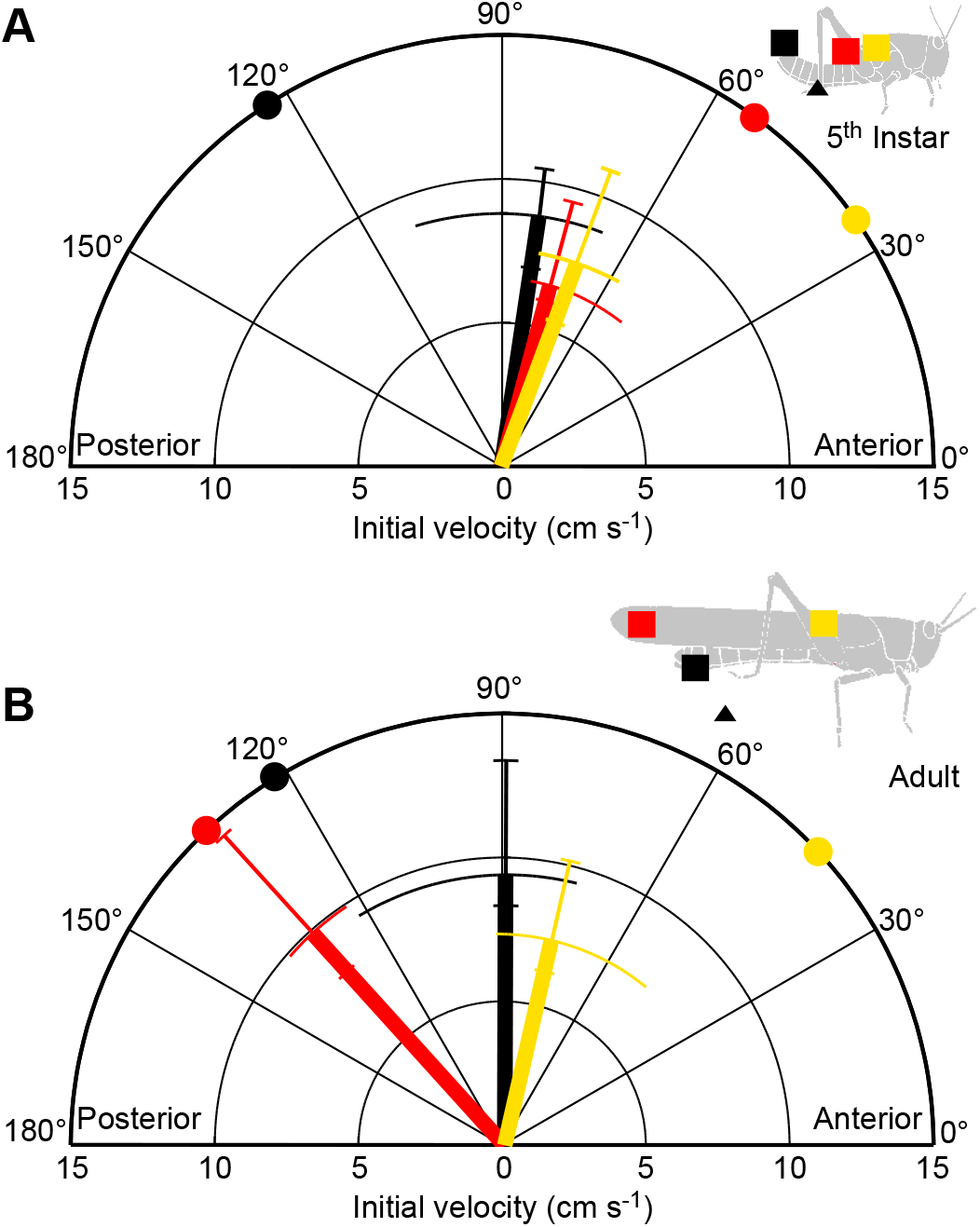
Initial movement trajectories of scratches elicited by touch of wing tip sites differ in 5^th^ instars and adults. The initial movement vector (direction and speed) of the distal tibia was measured over the first 200 ms for movements in 5^th^ instars (A) and adults (B), in response to touches at wing base (yellow), wing tip (red) and abdomen tip (black) targets. The median direction is indicated by the angle of each thick line, and the interquartile range is indicated by the length of the perpendicular thin curved error bars. The median speed is indicated by the length of each thick line and the interquartile range by corresponding thin error bars. The ‘ideal’ vector angle for each target is indicated by a colour-coded circle on the perimeter of the polar plot. The start and target locations are also indicated in the inset diagrams of 5^th^ instar and adult locusts. Movements toward wing tip targets were directed more posteriorly in adults than in 5^th^ instars. Speed was similar for 5^th^ instars and adults for movements towards wing base and abdomen tip, but different for movements towards wing tip.

For adults, initial movements toward wing tip targets (red in Fig. 4B) were directed more posteriorly than those for either wing base (yellow) or abdomen tip (black) targets (Fig. 4B). Target position influenced the initial movement direction (Friedman *χ*^2^_2,6_ = 9.000, *P* = 0.011).

There was no evidence for a difference between 5^th^ instars and adults in initial direction of movement towards wing base targets (yellow in Fig. 4A, B; Mann-Whitney *U* = 18, *P* = 1.000), or towards abdomen tip targets (black in Fig. 4A, B; Mann-Whitney *U* = 15, *P* = 0.631). The initial movement towards wing tip targets, however, was directed more posteriorly in adults than in 5^th^ instars (compare red vector directions in Fig. 4A and B; Mann-Whitney *U* = 0, *P* = 0.004).

For both 5^th^ instars and adults, the initial speeds of movement (vector lengths in Fig. 4A, B) were similar for scratches towards all three targets (Friedman Test, 5^th^ instars: *χ*^2^_2, 6_ = 3.000, *P* > 0.223; adults: *χ*^2^_2, 6_ = 4.333, *P* > 0.115).

There was no evidence for a difference between 5^th^ instars and adults in initial speed of movement towards wing base targets (yellow in Fig. 4A, B; Mann-Whitney *U* = 16, *P* = 0.749), or towards abdomen tip targets (black in Fig. 4A, B; Mann-Whitney *U* = 15, *P* = 0.631). There was weak evidence, however, that initial movement towards wing tip targets was slower in 5^th^ instars than in adults (compare red vector lengths in Fig. 4A and B; Mann-Whitney *U* = 5, *P* = 0.037). Overall these results indicate that movements elicited by touch of wing tip targets are aimed from their outset either anteriorly in 5^th^ instars or posteriorly in adults, suggesting different pre-determined trajectories.

### Cyclic component of scratching is influenced by target site

Most scratches consisted of an initial outward trajectory followed by several cyclical ‘loops’ near the target. Relatively few scratches (9.6 ±1.3 %, mean ± SEM) had no cyclical component (‘loops’). There was no difference in number of loops between 5^th^ instars and adults for movements towards wing base stimuli (χ^2^ = 1.201, df = 3, *P* = 0.867), abdomen tip stimuli (χ^2^ = 0.671, df = 3, *P* < 0.001) or wing tip stimuli (χ^2^ = 2.237, df = 3, *P* = 0.588).

The pattern of movement during the cyclic part of scratching was summarised as an effector probability distribution (Fig. 5). Regions of highest probability are darkest in Fig. 5, and the weighted centre of the distribution (‘centre of density’) is indicated by a circle (mean ± SEM).

**Figure 5.**
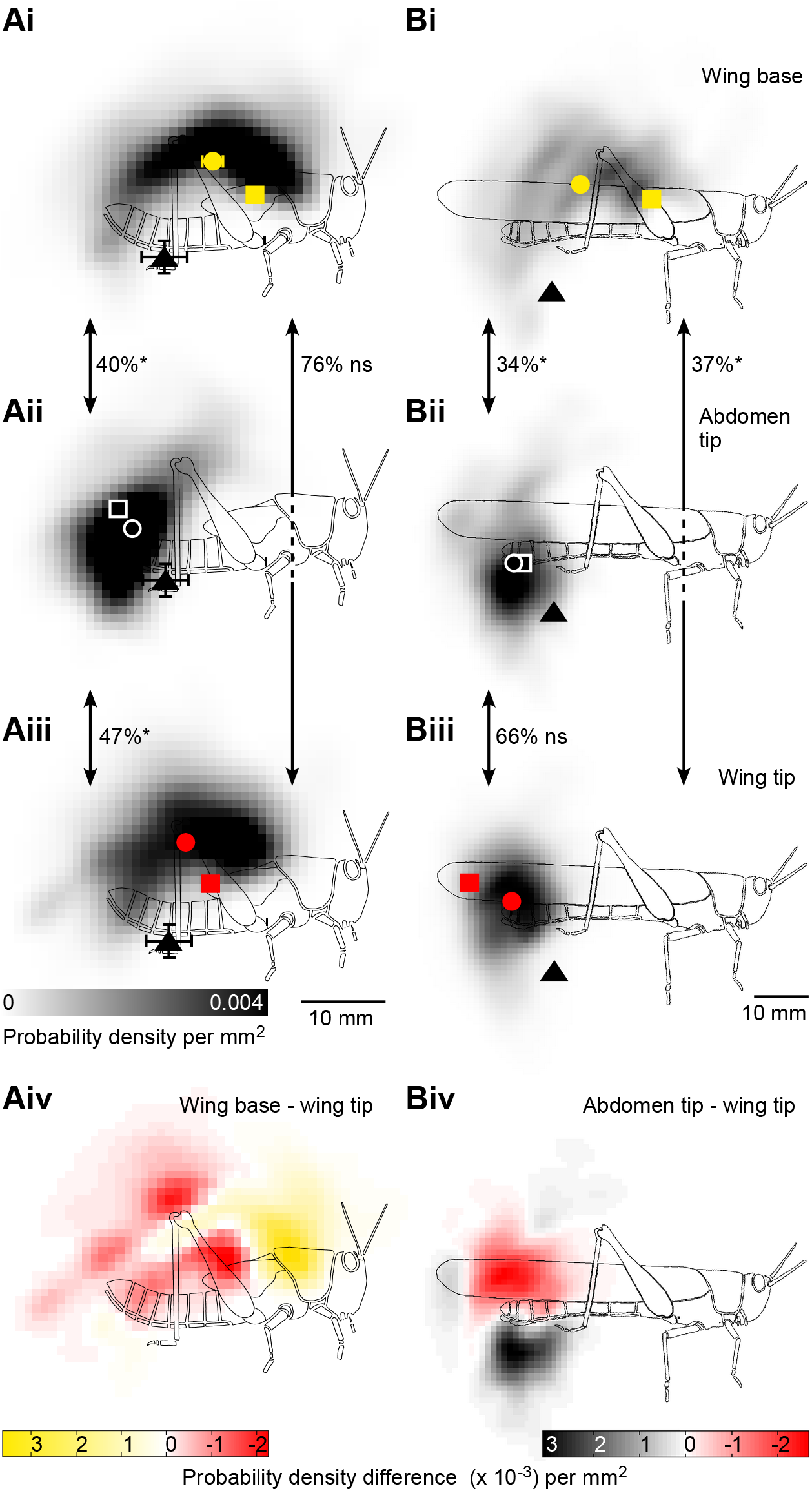
The cyclical components of scratches elicited by touch of wing tip sites differ in 5^th^ instars and adults. Grey scale probability distributions indicate the likelihood with which a particular point in the workspace was crossed by the distal tibia for 5^th^ instars (Ai-iv) and adults (Bi-iv), during movements elicited by touches at wing base (Ai, Bi), abdomen tip (Aii, Bii) and wing tip (Aiii, Biii) targets (colour coded squares). The start locations are indicated by black triangles (mean + SEM, most error bars within the symbol). The centre of density of each probability distribution is represented by a colour coded circle (mean ± SEM, errors bars for adults lie within the symbols). The overlap between pairs of distributions (double-headed arrows) is indicated by percentage values, and the significance (see Methods) by asterisks (significant) or ‘ns’ (not significant). For 5^th^ instars, the movement distribution for wing base targets (Ai) was biased anteriorly compared to that for wing tip targets (Aiii), as emphasised by the colour coded distribution of differences in probability (Aiv, red shading indicates areas where probability is higher for wing tip targets, yellow for wing base targets). Similarly, for adults the movement distribution for abdomen tip targets (Bii, black coding in Biv) was biased ventrally compared to that for wing tip targets (Biii, red coding in Biv). The number of scratches (from six 5^th^ instars and six adults) comprising each distribution in Ai-Biii are: 5^th^ instar wing base, n = 93; abdomen tip, 84; wing tip, 86; adult wing base, 83; abdomen tip, 80; wing tip, 79. The ‘number of frames required’ / ‘mean number of frames’ (see Methods) for each comparison indicated by double headed arrows is: 5^th^ instar wing base *vs* abdomen tip, 15 / 37; wing base *vs* wing tip, 223 / 49; abdomen tip *vs* wing tip, 23 / 36; adult wing base *vs* abdomen tip, 11 / 42; wing base *vs* wing tip, 11 / 49; abdomen tip *vs* wing tip, 107 / 37.

In 5^th^ instars, the probability distributions for movements aimed at targets at the base of the wing or the tip of the wing both formed arcs (shaded regions in Fig. 5Ai, Aiii) that curved from near the start positions (black triangles), dorsally and anteriorly, then ventrally and further anteriorly to near the targets (squares). The mean centres of density yellow and red circles respectively) were dorsal and posterior to the targets in both cases, so that the targets lay near the anterior ventral margin of the probability distributions. The probability distribution for movements aimed at targets at the tip of the abdomen formed a compact distribution over the target site (Fig. 5Aii). The centre of density (white circle) was close to the target (white square).

Probability distributions for movements aimed at the wing base and abdomen tip in 5^th^ instars overlapped by 40% but could be reliably distinguished (Fig. 5Ai-ii), as could those for movements toward wing tip and abdomen tip targets (47% overlap; Fig5Aii-iii). These movement distributions were thus target-specific. Probability distributions for movements aimed at the 5^th^ instar wing base and wing tip targets (which are separated by only 4 mm) overlapped by 76% and could not be reliably distinguished (Fig. 5Ai, Aiii). The distribution for the more anterior wing base target was nevertheless biased anteriorly compared to that for the more posterior wing tip target (Fig. 5Aiv).

Target position (wing base, wing tip or abdomen tip) had a strong effect on the anterior-posterior (x-axis) location of the centre of density of the probability distribution (Repeated Measures ANOVA *F*_2, 10_ = 31.13 *P* < 0.001). There was no evidence for a difference between the x-axis locations of the centres of density for movements aimed at wing base and wing tip targets (post-hoc *P* = 0.110). Both were, however, more anterior than the centre of density for movements towards abdomen tip targets (wing base: *P* = 0.004, wing tip: *P* = 0.004).

In adults, the probability distribution for movements aimed at targets on the base of the wing formed an arc that curved from near the start position (black triangle in Fig. 5Bi), dorsally and anteriorly, then ventrally and further anteriorly to near the target (square). The mean centre of density (circle) was posterior to the target. In adults, the probability distributions for movements aimed at targets at the abdomen tip or wing tip formed compact clusters close to the corresponding target sites (squares in Fig. 5Bii, Biii). The centre of density (circle) was ventral and anterior to the target (square) in movements towards the wing tip (Fig. 5Biii), whereas in movements towards targets at the abdomen tip the centre of density overlapped the target site (Fig. 5Bii).

To test whether movements were target-specific in adults, the overlap between pairs of probability distributions was compared in the same way as for 5^th^ instars. The probability distributions for movements aimed at wing base and wing tip targets overlapped by only 37% and could be distinguished reliably (Fig. 5Bi, Biii), as could movements aimed at the wing base or abdomen tip (34% overlap, Fig. 5Bi-ii). Probability distributions for movements aimed at wing tip and abdomen tip targets overlapped by 66% and could not be distinguished reliably (Fig. 5Bii-iii). The distribution for the more dorsal wing tip target was nevertheless biased dorsally compared to that for the more ventral abdomen tip target (Fig. 5Biv).

As for 5^th^ instars, target position had a highly significant effect on the x-axis location of the centre of density (Repeated Measures ANOVA, *F*_2, 10_ = 61.05, *P* < 0.001). In adults, movements towards the wing base had a more anterior centre of density than movements towards either the abdomen tip (post-hoc *P* = 0.002) or the wing tip (*P* = 0.001). In contrast movements towards the abdomen tip and wing tip had centres of density in a similar location along the x-axis (*P* = 1.000). Overall, these results indicate that both 5^th^ instars and adults accurately aim the cyclical part of their scratching movements at the correct target sites.

### Comparison of 5^th^ instar and adult scratching movements

For movements towards an anterior target such as the base of the wing, the probability distributions for 5^th^ instars and adults both had a similar shape and similar centres of density (Fig. 5Ai, Bi). The adult distribution was less dense because the trajectories covered a larger area (note scale bars in Fig. 5). For movements towards a posterior target, such as the tip of the abdomen, the probability distributions for 5^th^ instars and adults also had similar shapes and centres of density (Fig. 5Aii, Bii). The location of the wing tip changes from an anterior position in 5^th^ instar locusts to a posterior position in adults, and this was accompanied by a clear change in the behavioural response. In response to stimulation of the wing tip, 5^th^ instars made an anteriorly directed movement (Fig. 5Aiii) that had a similar shape to the probability distribution for movements aimed at the (anterior) base of the wing (Fig. 5Ai), whereas adults made a posteriorly directed movement (Fig. 5Biii), that had a similar shape to the probability distribution for movements aimed at the (posterior) tip of the abdomen (Fig. 5Bii).

### Joint kinematics are similar for corresponding targets in 5^th^ instars and adults

To determine whether 5^th^ instars and adults use similar patterns of inter-joint coordination to reach anterior or posterior targets, the combined thoraco-coxal + coxo-trochanteral joint angle was plotted against the femoro-tibial joint angle at the closest point of approach to the target (Fig. 6). Mean joint angles for each animal (coloured circles in Fig. 6) were superimposed on a large dataset of individual adult scratches from Dürr and Matheson (2003) (grey circles in Fig. 6). Dürr and Matheson (2003) stimulated multiple sites along the length of the adult wing surface and showed that combinations of joint angles were gradually shifted as the target position was moved along the wing’s length. Adult data in the present study correspond well with this pattern: for anterior wing base targets the thoraco-coxal + coxo-trochanteral joint angle was large and the femoro-tibial joint angle small (yellow circles in Fig. 6A), reflecting elevation of the femur and flexion of the tibia at the point of closest approach to the target. For posterior wing tip targets, the thoraco-coxal + coxo-trochanteral joint angle was large, and the femoro-tibial joint angle small (red circles in Fig. 6A) reflecting depression of the femur and extension of the tibia at the point of closest approach to the target. Joint angle combinations used to reach the abdomen tip lay on the same curve and were near those used to reach the wing (black circles in Fig. 6A).

**Figure 6.**
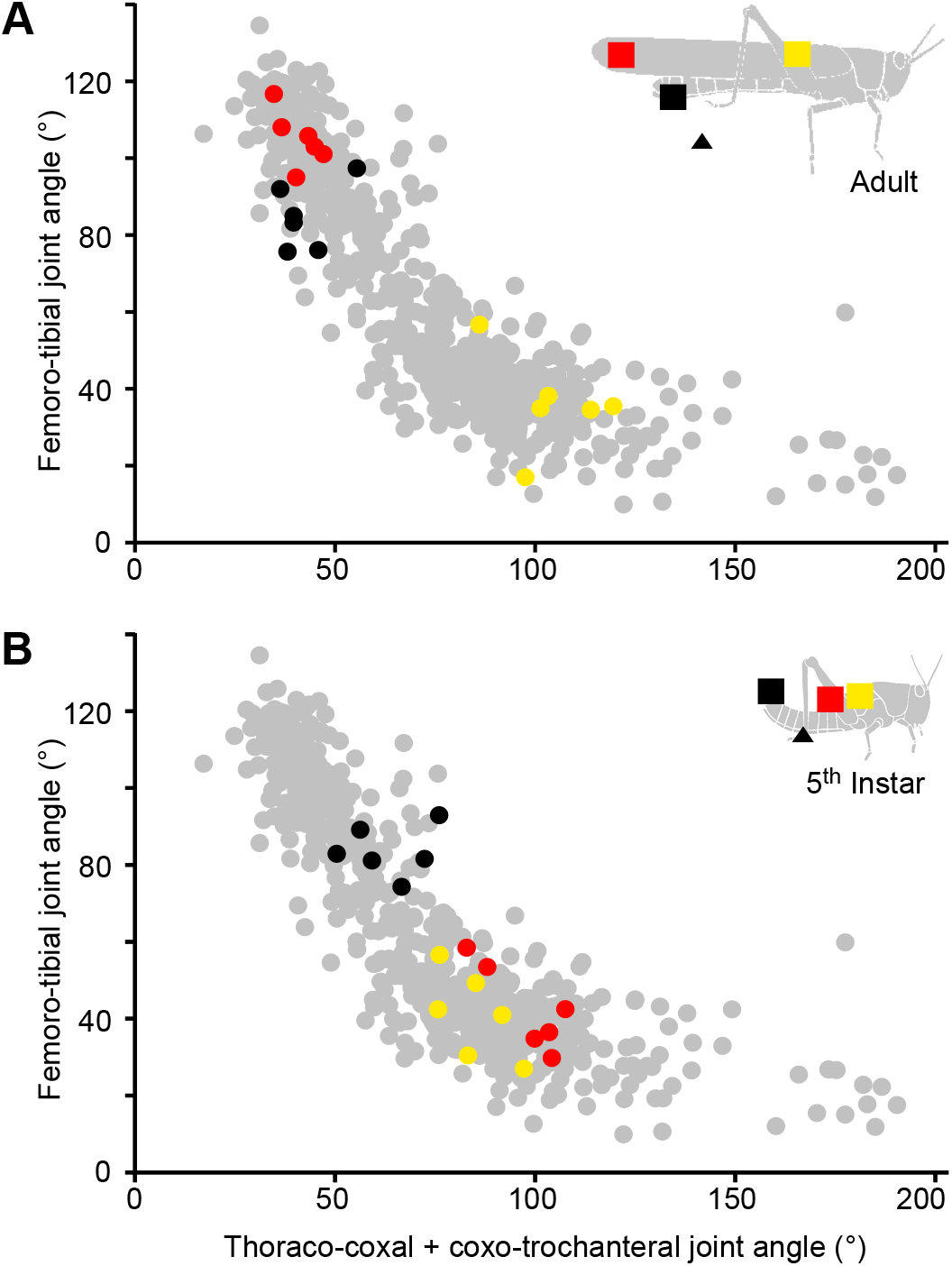
Hind leg joint angles at the point of closest approach to the wing tip target differ in 5^th^ instars and adults. Grey circles show combinations of femoro-tibial angle and the combined thoraco-coxal + coxo-trochanteral angle of the hind leg at the point of closest approach to the wing base for individual scratches made by adult locusts (data from Durr and Matheson, 2003). Superimposed on these are the mean values for the six adult (A) and six 5^th^ instar (B) locusts in the present study. The start (triangles) and target (squares) locations are colour coded in the inset diagrams of 5^th^ instar and adult locusts. For both 5^th^ instar and adult locusts, hind leg joint angles at the closest point of approach to each target fall on a continuum. In this joint angle space, movements to wing tip targets (red) lie near wing base targets (yellow) for 5^th^ instars (B), but near abdomen tip targets (black) for adults (A).

For 5^th^ instars, movements towards the anterior wing base target used joint angle combinations that coincided with those used to reach the same anterior target in adults (yellow circles in Fig. 6B *cf.* 6A). The combinations of joint angles used to reach the posterior abdomen tip target in 5^th^ instars similarly coincided with those for the abdomen tip adults (black circles in Fig. 6B *cf*. 6A). In contrast, the combinations of joint angles for movements aimed towards the wing tip were different in 5^th^ instars and adults (red circles in Fig. 6B *cf*. 6A), while still lying on the same curve. Specifically, the combinations of joint angles used to reach the wing tip target in 5^th^ instars were similar to those used to reach the wing base targets in both adults and 5^th^ instars, whereas those used to reach the wing tip target in adults were similar to those used to reach the abdomen tip targets of both adults and 5^th^ instars. Fifth instars used systematically larger femoro-tibial joint angles for a given thoraco-coxal + coxo-trochanteral angle to reach wing tip targets than they did for wing base targets (red circles in Fig 6B lie above yellow circles).

## DISCUSSION

We show that gentle tactile stimulation of the wings elicits aimed hind leg movements in juvenile locusts that closely resemble those of adults. Despite marked allometric growth and radical changes in wing morphology during imaginal development, these aimed movements are immediately retargeted to appropriate locations, while the characteristic form of the movements remains the same.

### Fifth instar and adult locusts use the same effector and movement patterns

Fifth instar locusts, like adults (Berkowitz and Laurent, 1996, Matheson, 1997a,b; Dürr and Matheson, 2001) perform aimed hind leg scratching movements in response to, and accurately aimed at, stimuli on the wings and abdomen. First, both 5^th^ instars and adults use the distal tibia as the effector. Second, both move the effector at similar speeds for the initial 200 ms of movement. Third, both achieve a similar accuracy, as determined by the minimum distance between the effector and the target. Fourth, both use comparable combinations of hind leg joint angles to reach a range of targets. A single movement pattern that can be shifted towards anterior or posterior targets on the wings (Dürr and Matheson, 2001) thus appears to be conserved across the imaginal moult. Combinations of joint angles lie on a continuum in joint angle space, allowing the effector to reach anterior targets using joint angle combinations at one end of the continuum (thoraco-coxal + coxo-trochanteral joint rotated anteriorly, and the femoro-tibial joint fully flexed), whereas posterior targets are reached using joint angle combinations at the other extreme of the continuum (thoraco-coxal + coxo-trochanteral joints rotated posteriorly, and the femoro-tibial joint extended). Targets between these two extremes are reached by intermediate joint angle combinations in both 5^th^ instars and adults. We therefore hypothesise that essentially the same central neuronal network, with properties conserved across the imaginal moult, generates scratching in 5^th^ instars and adults, but that the sensory inputs eliciting the behaviour must be modified.

### Isometric growth can explain the maintenance of aiming accuracy for targets on the abdomen

Isometric growth of the hind legs means that combinations of joint angles used by 5^th^ instars to reach locations on the abdomen will continue to direct the larger adult hind leg to the same region of the larger adult abdomen. Nevertheless, growth must also lead to changes in muscle mass and strength, leg mass, cuticular properties and joint mechanics (Andersen, 1973, 1974; Norman, 1995), so it remains possible that leg motor patterns used by 5^th^ instars differ from those used by adults to achieve the same kinematics.

Praying mantises use rapid aimed lunges of their fore legs and body to grasp moving prey. Striking distance is precisely estimated by a binocular triangulation mechanism of the visual system (Maldonado and Levin, 1967), with the longest distance that elicits a strike being related to the length of the fore legs in each developmental stage (Balderrama and Maldonado, 1973). The growth of the head and the eyes is such that throughout development the fovea registers a stimulus at the maximum catching distance. This triggers a striking movement at a distance that is appropriate for the length of the limbs at each developmental stage (Maldonado *et al.* 1974). In an analogous way, as the locust abdominal body wall elongates during development, distal targets on it move posteriorly, but it is likely that little or no remapping of sensory neurons is required, because unchanged patterns of connectivity with motor networks would ‘automatically’ drive the isometrically larger hind leg to the new target location, at least within the levels of accuracy seen for these movements. Such automatic compensation is not possible for wing tip targets, however, because of allometric wing growth and wing rotation.

### Developmental wing rotation at the imaginal moult does not explain changed responses to wing tip stimulation

Touching the exposed fore- or hind wing tips of 5^th^ instar locusts elicits anterior movements whereas touching the wing tips of adults elicits posterior movements. We have ruled out the possibility that 5^th^ instars fail to make posterior movements simply because the receptors are hidden under the (rotated) wing buds: stimulating the normally hidden dorsal fore wing receptors in 5^th^ instar locusts leads to *anterior* movements. The central circuits underlying posterior scratching are present in 5^th^ instars – since distal abdominal stimulation elicits posteriorly directed movements – but they do not produce posterior movements in response to wing tip stimulation until adulthood.

### Remodelling of sensory inputs may explain changed responses to wing tip stimulation

In contrast to the apparently learned ability of adult locusts to spontaneously walk across larger gaps than juveniles (Ben-Nun *et al.* 2013), appropriately aimed scratching movements did not require prior experience of either stimulation or movement following the adult moult. The most parsimonious explanation of the observed change in aiming is that the adult fore wing develops new tactile hair receptors that drive the existing central motor circuits to produce more posteriorly directed movements than those driven by comparable receptors on the juvenile wing buds. Aimed scratches are elicited by activation of touch sensitive hairs distributed across the wing surface, with a dorsally-directed row on the adult forewing postcubitus vein being particularly efficacious (Page and Matheson, 2004). The somatosensory representation of the wing surface is unknown for any insect – in part because the afferent neurons appear to have small diameters, yet long axons, which makes staining difficult. It is reasonable to assume, however, that the afferents form a somatotopic map in the local ganglion, as is the case for locust leg and body wall receptors (Newland, 1991; Newland *et al.* 2000). We propose that the juvenile wing bud somatosensory map, and its connections to leg motor networks, is extended ‘posteriorly’ through the addition of new receptors born on the adult wing during the final larval instar (Altman *et al.* 1978). In the cricket, clavate mechanosensory hairs and their receptors are likewise added at each moult. The central projections of later-born receptors project less far, and less extensively, into the terminal abdominal ganglion, so that the population of afferents forms a topographic representation of the surface of the cercus depending on both birth date and hair location (Murphey *et al.* 1980; Murphey, 1981). The number of wind-sensitive hairs on the head of a locust similarly increases with each instar (Sviderskii, 1969), and this is accompanied by a progressive development of the flight motor pattern – which they drive – in juveniles and adults (Altman, 1975; Kutsch, 1971, 1985). Immobilising the wings at the imaginal moult does not prevent expression of the adult flight motor pattern later in adult life (Altman, 1975), and the development of the final adult wingbeat frequency does not require flight experience, suggesting that, as for aimed scratching, changes are ‘hard-wired’ rather than learned. Although the flight motor pattern begins to develop during larval stages, the full adult pattern is not expressed until the day of (following) the imaginal moult. We have shown that although the motor system of 5^th^ instar locusts can produce posteriorly aimed hind limb movements (e.g. in response to touch of the abdomen), such posterior movements are not elicited by touch of the wings until the adult wings are fully expanded following the imaginal moult, when the new stimulus-response relationship is apparent even on the very first touch of the new sensory surface. Locusts thus possess a mechanism to automatically compensate their aimed limb movements for allometric growth of the wing sensory surface.

## MATERIALS AND METHODS

### Animals and experimental protocol

#### Movement kinematics

Experiments were carried out on six 5^th^ instar (3 male, 3 female) and six adult (3 male, 3 female) desert locusts *(Schistocerca gregaria* Forskål) taken from a 556laboratory culture established at the University of Leicester, UK with stock from Blades Biological Ltd. (Kent, UK) and the Department of Zoology, University of Cambridge, UK. Animals were maintained in crowded cages under a 12: 12 h light: dark photoperiod with a corresponding 36°C: 25°C thermal regimen, and fed *ad libitum* with fresh wheat seedlings and organic wheat bran. For experiments, animals were tethered with a fine loop of wire that passed around their pronotum without obstructing movements of their legs. They were suspended above a foam ball on which they could stand or walk. The tether allowed each animal to adjust its body posture, with the exception of thorax height above the substratum which was set to the height normally maintained by a walking locust. The hind leg was manipulated into a standardised start posture (black triangles in Fig. 7Ai, Bi) by placing the tarsus on a horizontal rod located at 67% of the distance between the anterior rim of the metathoracic coxal joint and the tip of the abdomen (Dürr and Matheson, 2003). The eyes and ocelli were covered with black acrylic paint (Cryla, Daler-Rowney, Bracknell, UK) to exclude visual cues. Experiments were carried out at 22-24°C, maintained by an infra-red heat lamp suspended 50cm above the animal. To facilitate video based motion tracking, 1 mm diameter discs cut from reflective tape (3M Scotchlite), were attached to the body and the right hind leg with insect glue (Thorne (Beehives) Ltd., UK; Fig. 7Aii, Bii). Movement of the unguis was ignored. For illustrative purposes tarsal length was standardised to 6.75 mm. See Dürr and Matheson (2003) for additional details.

**Figure 7.**
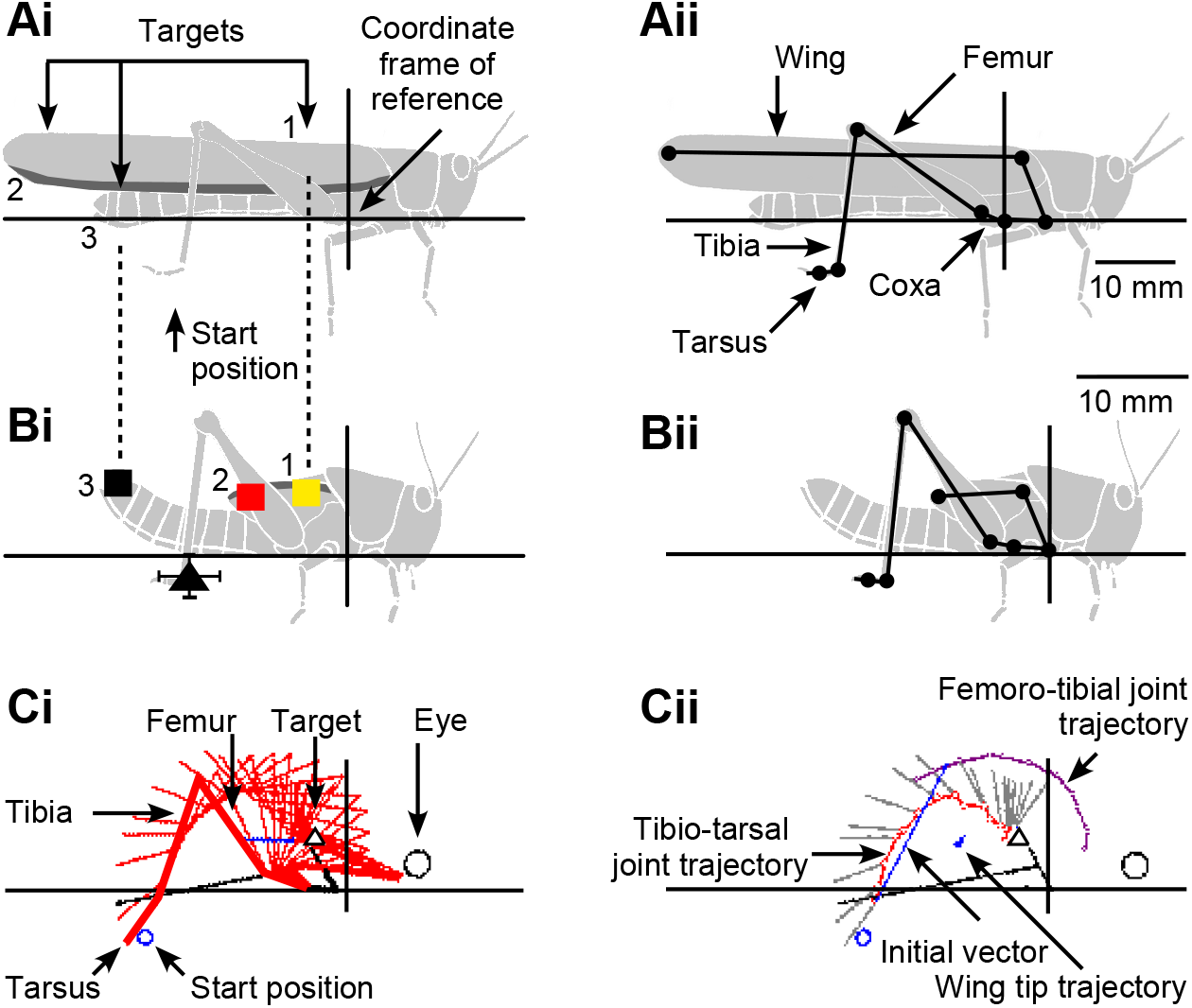
Coordinate frame of reference, start and stimulus (‘target’) positions for adults (Ai) and 5^th^ instars (Bi). Squares indicate mean stimulus locations on the wing base (yellow), wing tip (red) or abdomen tip (black). Errors (± SEM) lie within the symbol for all targets (N = 6 5^th^ instars and 6 adults). The hind leg tarsus was placed on a rod at a single start position (black triangle, mean ± SEM). The body-centred coordinate frame of reference had its origin on the anterior rim of the hind leg coxa for adults (black axes in Ai) and was shifted so that wing base and abdomen tip stimuli in 5^th^ instars were aligned with those of the adults (broken lines). Movements were tracked using reflective markers placed on the body and limbs (black circles in Aii, Bii) and fitted with a body model (black lines joining circles in Aii, Bii). For a single scratch it was thus possible to visualise in each video frame the locations of the hind leg segments (Ci, red lines) and wing midline (blue line) relative to the thoracic frame of reference (short black lines). The abdomen (long black line, partly obscured) was marked in the first frame of each sequence only, but did not move during a scratch. The trajectories (Cii) of the hind leg tibio-tarsal joint (red dots and line) and femoro-tibial joint (purple dots and line) are shown for a wing base stimulus (open triangle) in a 5^th^ instar locust. Grey lines show the tarsus position in each frame (20 ms between frames), and the blue line indicates the initial movement vector for the first 200 ms of movement.

Scratching movements of the right hind leg ipsilateral to stimulus sites on a wing were analysed in the following way. The adult fore wing was notionally divided into 5 regions of equal length. The most anterior and posterior of these regions were used as ‘wing base’ and ‘wing tip’ stimulus sites, respectively, and the posterior fifth of the abdomen was used as a posterior ‘abdomen tip’ target (Fig. 7Ai). For 5^th^ instar locusts, the wing pad was divided into anterior ‘wing base’ and posterior ‘wing tip’ regions (Fig. 7Bi). Stimulation was provided by gently touching the wing base, wing tip or abdomen tip with a fine paint brush to elicit scratching behaviour (mean ± SEM stimulus positions are indicated by coloured squares in Fig. 7Ai, Bi). Wing base, wing tip and abdomen tip stimuli were given randomly until at least 10 scratches were recorded for each stimulus site. The main results are based on analyses of 793 scratches in 12 locusts. Sample sizes were guided by our published data (e.g. Dürr and Matheson, 2003).

Locusts rarely touched their own body surface during the first few cycles of a scratching response, so responses were open-loop with respect to the stimulus. Responses in which the leg made contact with the body surface or with the stimulating brush were analysed up until the time of first contact. Contact was determined by visual inspection of the video.

#### Responses to dorsal fore wing stimulation in 5^th^ instars

Ten female and two male 5^th^ instar animals were used to examine responses to touch of the normally hidden dorsal fore wing surface. Animals were tethered as described above and their eyes covered with black paint. They walked freely on the ball and the hind leg was not placed onto a standard starting rod. The left hind wing bud and both of the wing buds on the right were ablated using fine scissors, leaving only short stumps of approximately ¼ of the total length of each wing bud. The inner (morphologically dorsal) surface of the remaining intact left fore wing bud was then stimulated with a fine paintbrush. Each animal received 10 stimuli at 1 min intervals. All but one female animal responded to at least one stimulus with an aimed scratching movement (mean 3.4 responses per animal, median 3.5 responses per animal). The non-responding animal was omitted from further analysis. The responses were scored as: *Anterior*, if the femur was strongly rotated anteriorly so that the tarsus passed anterior to the midpoint of the abdomen (which corresponds approximately to the tip of the wing bud, see Fig. 1Bi); *Middle*, if the femur rotated anteriorly so that the tarsus passed between the tip and the midpoint of the abdomen; and *Posterior*, if the femur was rotated posteriorly so that the tarsus passed posterior to the tip of the abdomen (as it would for a scratch aimed at the abdomen tip or adult wing tip (see Fig. 1Bi, ii). Trials that elicited no scratching response were scored as *None*. Each animal was also tested with a touch to the tip of the abdomen, which in all cases led to robust *Posterior* scratches.

#### Development of aimed movements following the imaginal moult

Nine animals were observed moulting from 5^th^ instar to adult in their normal home cage. A fine paintbrush was used to touch the dorsal distal fore wing tip and other locations on the wings and body. Animals were observed continuously from the point at which the old exoskeleton split, and were stimulated periodically until the wings were fully expanded and rotated into their adult orientation (30-60 min). After emergence from the old exoskeleton, animals hung from a vertical support, head up, while the adult wings first expanded in their juvenile orientations (i.e. with the hind wing outermost, and both wings rotated so that their dorsal surfaces faced the body). During this period animals remained in the same posture with little movement unless disturbed. In a brief movement, animals then rotated and adjusted the expanded wings so that they assumed their normal adult orientations. Responses to all stimulations were recorded.

A further 13 animals (6 male, 7 female) were also observed moulting, and had both hind legs restrained within 1 min after emergence from the old exoskeleton – before the animal had a chance to make any aimed scratching movements and before the cuticle had hardened. A loop of 370μm diameter tinned copper wire was passed over the femoro-tibial joint of each fully flexed hind leg to restrain the tibia in the fully flexed position. The limb was free to move at the thoraco-coxal and coxo-trochanteral joints, but the animal could not extend the tibia or make aimed scratching movements. Animals were maintained together for 20-24 h in small holding cages (25 cm L × 15cm W × 20 cm H) in which they could walk, climb and feed. For testing, animals were suspended above a foam ball as described in *Kinematic analyses* above, with their eyes blacked out. They were not provided with a start rod. The wire loop was cut with fine wire-cutters to free the tibia and then the fore wing tip was immediately stimulated with a fine paintbrush. Responses were observed and recorded as either ‘walking/struggling’ or as ‘anterior’, ‘middle’ or ‘posterior’ scratches as defined in *Responses to dorsal fore wing stimulation in 5^th^ instars* above. Subsequent stimulation of the fore wing base and tip permitted repeated observations.

### Video acquisition and analysis

For kinematic analyses, animals were videotaped using a colour CCD camera (C1380, JVC, Yokohama, Japan) operated at a shutter speed of 1/500 s. The sVHS video signals were combined on a multiviewer (MV-40PS, For-A, Japan) and time stamped using a video timer (VTG-33, For-A, Tokyo, Japan). Images were taped on an sVHS video recorder (HR-S7500, JVC, Yokohama, Japan), displayed on a monitor (PVM-1450MD, Sony, Tokyo, Japan) and played back for capture by a personal computer video interface card (MiroVIDEO DC30 plus, Pinnacle Systems, California, USA) in PAL video format at a size of 720 × 540 pixels using MiroVIDEO Capture software (25 Hz frame rate for full interlaced video frames) and compressed using Microsoft “dvsd” codec. A deinterlacing algorithm (Videotrack 2D written by J. Zakotnik, available on request from the corresponding author) then split each frame into even and odd frames, and linearly interpolated the missing lines to give a final frame rate of 50 Hz. Short clips containing individual scratches were produced in VirtualDub (A. Lee, http://www.virtualdub.org/index). A calibration target was filmed during every experiment.

Videotrack 2D was used to detect the co-ordinates of the eight markers within each frame that described the animal’s body position and leg movement (Fig. 7 Aii, Bii. See Dürr and Matheson 2003 for a discussion of the accuracy of this method). The line between the markers adjacent to the hind leg coxa and the front leg coxa defined the x-axis of the body-centred coordinate frame, with the marker 2 mm in front of thoraco-coxal joint of the hind leg set to the origin (Fig. 7 Aii, Bii). The stimulus location and the start position of the tarsus were digitised on the first frame of each behavioural sequence. For the x-axis comparison of centres of density, the 5^th^ instar coordinates were scaled by 140%, based on the mean difference in size for six 5^th^ instar and six adult locusts, and wing base and abdomen tip stimulus locations were aligned by shifting the 5^th^ instar frame of reference anteriorly by 2.7 mm. This standardisation permitted the direct comparison of movement trajectories between 5 ^th^ instars and adults of different sizes.

### Data analysis

Analysis of individual scratching responses started with the frame before the movement began and ended if the tarsus touched the ground, the tarsus hit the brush, the leg completed three cycles of movement (‘loops’, see below), or the leg stopped moving for longer than 120 ms. The 2-dimensional coordinates of hind leg joints throughout a scratch were output by Videotrack 2D and subsequently analysed by Scratch Analysis (written by V. Dürr, available on request from the corresponding author; Fig. 7Ci,ii). The position of the hind leg relative to the body is shown as an example for one movement toward a wing base target in a 5^th^ instar locust (Fig. 7Ci). The hind leg tarsus moved from the start position (small blue circle) anteriorly and dorsally toward the stimulus position (triangle) on the wing (blue line) driven by anterior rotation of the thoraco-coxal joint and flexion of the femoro-tibial joint. Figure 7Cii shows the sequential locations of the femoro-tibial joint (purple), the tibio-tarsal joint (red), and the wing bud tip (blue) for the same movement. Scratch Analysis computed joint angles and angular velocities for all four hind leg joints in each frame of the sequence. Statistical tests were calculated with the Statistical Package for the Social Sciences (SPSS Version 12, SPSS Inc. Chicago, USA). For all data, apart from the density maps described below, means were first calculated for each animal and then averaged across animals and presented together with the standard error of the mean. The outcomes of statistical tests are described using the nuanced terminology recommended by Colquhoun (Colquhoun, 2014; Colquhoun, 2015): p > 0.1, ‘no evidence for a real effect’; 0.05 < p < 0.1, ‘very little evidence for a real effect’; 0.01 < p < 0.05, ‘weak evidence for a real effect’; 0.001 < p < 0.01, ‘moderate evidence for a real effect’; p < 0.001, ‘strong evidence for real effect’.

To determine which part of the hind leg was being aimed at the target, the hind leg was divided into 24 ‘Units’ of equal length (see Dürr and Matheson, 2003) and the shortest distance between each Unit and the target during the scratch was calculated. Having determined the point on the leg that on average most effectively reached all 3 target sites, the movement trajectory of this effector point (≈4 mm proximal to the tibio-tarsal joint) was used to define the closest point of approach to the target in each scratch, thus providing a measure of the accuracy of each movement. A mean accuracy was established for each animal for movements towards wing base, wing tip and abdomen targets.

Two components of each scratching response were analysed: a short (200 ms) initial vector component (Fig. 7Cii) and a second cyclic component (Dürr and Matheson, 2003). The first 200 ms of movement was used to calculate the initial speed and angle of movement of the effector (scaling was not applied to these values). The cyclic component was quantified by first determining the number of movements with 0, 1, 2, or 3 loops for 5^th^ instars and adults for each target. A loop was defined by both extension and flexion of the femoro-tibial joint, and ended with the transition of the femoro-tibial joint from flexion to extension. The cyclic component was further characterised by calculating a probability distribution of the location of the distal tibia across scratches (see Dürr and Matheson, 2003 for details). The centre of density of the probability distribution was used as a single measure of the location of the scratch. To distinguish between pairs of movement distributions the probability distributions were treated as empirical likelihood functions, and Bayes’ rule was applied to decide which one of the two sites had most likely been stimulated to cause the leg to move across point X during a single video frame (see Dürr and Matheson, 2003 for derivation and details). The likelihood of the probability estimate being wrong for a single observation (frame) was used to determine a critical number of frames required to permit two maps to be distinguished. If the average number of frames of the two maps being compared was more than the critical number of frames needed to distinguish between them, then the two distributions were considered statistically different. In other words, the distributions were considered significantly different if observation of a single scratch of average duration would permit a correct prediction of the observed probability distribution in 95% of cases (see Dürr and Matheson, 2003). Probability distributions were compared for pairs of target sites to determine: (1) the percentage volume overlap of the two distributions; and (2) the minimum number of observations needed to obtain a 95% chance of making a correct decision (>50% correct single assignments).

The point of closest approach was shown in joint angle space for comparison with data from Dürr and Matheson (2003). Measurements of the thoraco-coxal and the coxo-trochanteral joint angles of the hind leg were smoothed with a sliding window having a width of five frames, weighted by binomial coefficients.

## ACKNOWLEDGEMENTS

We thank Jean Liggins, Maurice Andrews and Donna Betts for technical support, and Benjamin Matheson for help with tickling some of the locusts. Brendan O’Connor and Jan Ache provided valuable comments on a draft of the manuscript.

## AUTHOR CONTRIBUTIONS

TM conceived, obtained funding for, and supervised the project. AP and TM designed, carried out and analysed experiments. AP and TM wrote the paper.

## SUPPLEMENTARY MATERIAL

Original video recordings of all trials analysed in this work will be made available on the University of Leicester Figshare repository.

## REFERENCES

Altman, J.S. Changes in the flight motor pattern during the development of the Australian Plague Locust, Chortoicetes terminifera. J. Comp. Physiol. A 97: 127 – 142.

Altman, J.S., Anselment, E. and Kutsch, W. (1978). Postembryonic development of an insect sensory system: ingrowth of axons from hindwing sense organs in *Locusta migratoria*. Proc. Roy. Soc. Lond. B 202: 497 – 516.

Andersen, S.O. (1973). Comparison between the sclerotization of adult and larval cuticle in *Schistocerca gregaria*. J. Insect Physiol. 19: 1603 – 1614.

Andersen, S.O. (1974). Cuticular sclerotization in larval and adult locusts, Schistocerca gregaria. J. Insect Physiol. 20: 1537 – 1552.

Balderrama, N. & Maldonado, H. (1973). Ontogeny of the behaviour in the praying mantis. J. Ins. Physiol. 19: 319 – 336.

Ben-Nun, A., Guershon, M. & Ayali, A. (2013). Self body-size perception in an insect. Naturwissenschaften 100: 479 – 484.

Berkowitz, A. & Laurent, G.J. (1996a). Central generation of grooming motor patterns and interlimb coordination in locusts. J. Neurosci. 16: 8079 – 8091.

Burnett, G.F. (1951). Observations on the life-history of the red locust, Nomadacris septemfasciata (Serv.) in the solitary phase. Bull. Ent. Res. 42: 473 – 490.

Campbell, J.I. (1961). The anatomy of the nervous system of the mesothorax of *Locusta migratoria migratorioides* R. & F. Proc. Zool. Soc. Lond. 137: 403 – 432.

Chiba, A., Shepherd, D. & Murphey, R.K. (1988). Synaptic rearrangement during postembryonic development in the cricket. Science 240: 901 – 905.

Colquhoun, D. (2014). An investigation of the false discovery rate and the misinterpretation of p-values. Open Science 1, 140216.

Colquhoun, D. (2015). Comment on evidence levels. Available from: http://rsos.royalsocietypublishing.org/content/1/3/140216#comment-1889100957 (Accessed 29/4/2019).

Dirsh, V.M. (1967). The post-embryonic ontogeny of Acridomorpha (Orthoptera). Eos: Revista Española de Entomología 43: 413 – 514.

Dürr, V. & Matheson, T. (2003). Graded limb targeting in an insect is caused by the shift of a single movement pattern. J. Neurophysiol. 90: 1754 – 1765.

Easter, S.S. (1983). Postnatal neurogenesis and changing connections. Trends Neurosci. 6: 53 – 56.

Ivanova, T.S. (1947). Wing root development in *Calliptamus italicus* L. Rep. USSR Acad. Sci. LVI: 885 – 887.

Khattar, N. (1972). A description of the adult and the nymphal stages of *Schizodactylus monstrosus* (Drury) (Orthoptera). J. Nat. Hist. 6: 589 – 600.

Konczak J. & Dichgans J. (1997). The development toward stereotypic arm kinematics during reaching in the first 3 years of life. Exp. Brain Res. 117: 346 – 354.

Konczak, J., Borutta, M. & Dichgans, J. (1997). The development of goal-directed reaching in infants. 2. Learning to produce task-adequate patterns of joint torque. Exp. Brain Res. 113:465 – 474.

Kral, K. & Poteser, M. (2009). Relationship between body size and spatial vision in the praying mantis - an ontogenetic study. J. Orthoptera Res. 18:153 – 158.

Kutsch, W. (1971). The development of the flight motor pattern in desert locust, Schistocerca gregaria. Z. vergl. Physiol. 74: 156 – 168.

Kutsch, W. (1985). Pre-imaginal flight motor pattern in Locusta. J. Ins. Physiol. 31: 581 – 586.

Maldonado, H. and Levin, L. (1967). Distance estimation and the monocular cleaning reflex in praying mantis. Z. vergl. Physiol. 56: 258 – 267.

Maldonado, H., Rodriguez, E. & Balderrama, N. (1974). How mantids gain insight into the new maximum catching distance after each ecdysis. J. Ins. Physiol. 20: 591 – 603.

Matheson, T. (1997). Hindleg targeting during scratching in the locust. J. Exp. Biol. 200: 93 – 100.

Matheson, T. (1998). Contralateral coordination and retargeting of limb movements during scratching in the locust. J. Exp. Biol. 201: 2021 – 2032.

Matheson, T. (2002). Metathoracic neurons integrating intersegmental sensory information in the locust. J. Comp. Neurol. 444: 95 – 114.

Matheson, T. and Dürr, V. (2003). Load compensation in targeted limb movements of an insect. J. Exp. Biol. 206: 3175 – 3186.

Murphey, R.K. (1981). The structure and development of a somatotopic map in crickets: the cercal afferent projection. Dev. Biol. 88: 236 – 246.

Murphey, R.K., Jacklet, A. & Schuster, L. (1980). A topographic map of sensory cell terminal arborizations in the cricket CNS: correlation with birthday and position in a sensory array. J. Comp. Neurol. 191: 53 – 64.

Newland, P.L. (1991). Morphology and somatotopic organisation of the central projections of afferents from tactile hairs on the hind leg of the locust. J. Comp. Neurol. 312: 493 – 508.

Newland, P.L., Rogers, S.M., Gaaboub, I and Matheson, T. (2000). Parallel somatotopic maps of gustatory and mechanosensory neurons in the central nervous system of an insect. J. Comp. Neurol. 425: 82 – 96.

Norman, A.P. (1995). Adaptive changes in locust kicking and jumping behaviour during development. J. Exp. Biol. 198: 1341 – 1350.

Page, K.L. & Matheson, T. (2004). Wing hair sensilla underlying aimed hindleg scratching of the locust. J. Exp. Biol. 207: 2691 – 2703.

Roonwal, M.L. (1940). Preliminary note on some directional changes among locusts and other Acrididae, and the importance of the third instar. Indian J. Entomol. 2: 137 – 144.

Roonwal, M.L. (1946). Studies in intraspecific variation. II. New rules governing the correlations between normal-and extra-moulting and directional reversal of the elytron-wing complex in the desert locust and other Acrididae (Orthoptera). Indian J. Entomol. 7: 77 – 84.

Roonwal, M.L. (1952). Further observations on directional changes in locusts and other short-horned grasshoppers (Insecta: Orthoptera: Acrididae), and the importance of the third instar. Proc. Nat. Inst. Sci. India 18: 207 – 215.

Sviderskii, V.L. (1969). Receptors of the forehead of Locusta migratoria. J. Evol. Biochem. Physiol. 5: 482 – 490. [Translated from Russian]

Uvarov, B. (1966) Grasshoppers and Locusts: A Handbook of General Acridology. Volume Anatomy, Physiology, Development, Phase Polymorphism, Introduction to Taxonomy. Cambridge: Published for the Anti-Locust Research Centre at the Cambridge University Press. 481

